# Rad21l1 cohesin subunit is dispensable for spermatogenesis but not oogenesis in zebrafish

**DOI:** 10.1101/2020.09.23.309591

**Authors:** Yana P. Blokhina, Michelle Frees, An Nguyen, Masuda Sharifi, Daniel B. Chu, Bruce W. Draper, Sean M. Burgess

**Affiliations:** Department of Molecular and Cellular Biology, University of California, Davis, California, United States of America; Integrative Genetics and Genomics Graduate Group, University of California, Davis, United States of America; Biochemistry, Molecular, Cellular, and Developmental Biology Graduate Group, University of California, Davis, United States of America

## Abstract

Meiosis produces haploid gametes that will give rise to the next diploid generation. Chromosome segregation errors occurring at one or both meiotic divisions result in aneuploidy, which can lead to miscarriages or birth defects in humans. During meiosis I, ring-shaped cohesin complexes play important roles to aid in the proper segregation of homologous chromosomes. While REC8 is a specialized meiosis-specific cohesin that functions to hold sister chromatids together, the role of its vertebrate-specific paralog, RAD21L, is poorly understood. Here we tested if Rad21l1, the zebrafish homolog of human and mouse RAD21L, is required for meiotic chromosome dynamics during oogenesis and spermatogenesis. We found that Rad21l1 is an abundant component of meiotic chromosomes where it localizes to both the chromosome axes and the transverse filament of the synaptonemal complex (SC). Knocking out *rad21l1* causes nearly the entire mutant population to develop as fertile males, suggesting the mutation triggers a sex reversal from female to male due to a failure in oocyte production. The *rad21l1*^*−/−*^ mutant males display normal fertility at sexual maturity. Sex reversal was partially suppressed in the absence of *tp53,* suggesting that the *rad21l1*^*−/−*^ mutation causes defects leading to a Tp53 dependent response, specifically in females. The *rad21l1*^*−/−*^*;tp53*^*−/−*^ double mutant females produced elevated rates of decomposing eggs and deformed offspring compared to *tp53*^*−/−*^ controls. This response, however, is not linked to a defect in repairing Spo11-induced double-strand breaks since deletion of Spo11 does not suppress the sex reversal phenotype. Overall, our data highlight an exceptional sexually dimorphic phenotype caused by knocking out a meiotic-specific cohesin subunit. We propose that Rad21l1 is required for maintaining the integrity of meiotic chromatin architecture during oogenesis.

**Author Summary:** A prominent symptom of age-linked reproductive decline in women is the increased rate of miscarriage and birth defects due to aneuploidy. Aneuploidy can arise when chromosomes fail to segregate properly during meiosis, the process of creating haploid gametes from a diploid germ cell. Oocyte progression normally arrests prior to anaphase I, after homologous chromosomes have formed crossovers, but before ovulation, which triggers the first round of segregation. This prolonged arrest makes oocytes especially vulnerable to degradation of meiotic chromosome structure and homolog connections over time. Cohesin complexes play a major role in maintaining the meiotic chromosome architecture. Here we assess the role of the vertebrate-specific Rad21l1 cohesin subunit in zebrafish. We find that while males appear mostly unaffected by loss of Rad21l1, oocyte production is massively compromised, leading to sex reversion to males. Sex reversion can be partially prevented in the absence of Tp53, demonstrating that loss or Rad21l1 leads to a Tp53-dependent response in oocytes. Strikingly, double mutant *rad21l1 tp53* females produce large numbers of poor quality eggs and malformed offspring. This demonstrates a cohesin-linked vulnerability in female meiosis not present in males and sheds light on a potential mechanism associated with the decline in female reproductive health.

## Introduction

Meiosis is the cellular process that forms haploid gametes and drives the inheritance of chromosomes from one generation to the next. Two rounds of chromosome segregation following one round of DNA replication function to deposit the correct number of chromosomes in each gamete. Errors in this process can result in aneuploidy, a leading cause of birth defects and miscarriages in women. The majority of these errors occur during oogenesis. As women age, the incidence of pregnancies with trisomic and monosomic embryos may exceed 50% [1,2]. The cellular mechanisms that lead to aneuploidy in oocytes are poorly understood, yet several lines of evidence point to the premature degradation of cohesin complexes with age [3–8].

Cohesins are multi-subunit ring-like complexes that link two double-stranded DNA (dsDNA) strands together [9–11]. The complexes are composed of two SMC proteins (structural maintenance of chromatin), which interact to form the ring, and a kleisin subunit that functions to close the ring [12]. Combinations of different SMC and kleisin paralogs carry out a number of cellular functions, one of which is to maintain connections between sister chromatids during meiosis [13–17]. Sister chromatid cohesion, in combination with at least one crossover between homologous chromosomes, is essential to keep homologous chromosomes physically linked until they separate at anaphase I [15,18–21].

REC8 and RAD21L are two meiosis-specific kleisin subunits [12]. REC8 plays critical roles in forming and maintaining the unique chromosome architecture that supports the pairing, synapsis, and crossing over between homologous chromosomes and is conserved from yeast, plants, worms, flies, to mammals [13–15,18,21–30]. By contrast, RAD21L is only found in vertebrates. Several studies have linked RAD21L to a role in establishing interactions between homologous chromosomes in mouse spermatocytes [31–36]. The loss of *Rad21l* in mouse leads to infertility in the male, and age-dependent sterility in the female [32]. Understanding the role RAD21L plays in meiotic chromosome dynamics, however, has been elusive due to functional redundancy seen during the analysis of mutant phenotypes in mouse [37]. Although homologs of RAD21L have been identified in other vertebrate genomes, there is a dearth of studies of this cohesin in vertebrates other than mouse.

Zebrafish has emerged as an excellent model to use genetic approaches to study the chromosome events in meiosis [38–41]. Both sexes produce gametes throughout their lives, thus providing a window to study sexually dimorphic features of female and male meiosis. Importantly, hundreds of eggs from individual animals can be analyzed in a single cross, which provides a quantitative measure of gamete quality. The development of progeny is easily assessed, since embryos are transparent and develop outside the body. Lab strains of zebrafish do not have a heterogametic sex with unpaired or partially paired sex chromosomes as seen in many other vertebrate species. This bypasses some of the potentially confounding effects of disrupting meiotic sex chromosome inactivation (MSCI) that can lead to prophase arrest when homolog pairing is compromised [42,43]. For example, the prophase arrest phenotype seen in male *Rad21l1* mutant mice has been attributed to the unpaired X-Y sex body [31,32,37,44].

*Rad21l1* is the zebrafish homolog of mouse and human *Rad21L*. The aim of this study is to use zebrafish as a novel model vertebrate organism to study sex-specific roles of Rad21l1. We show that Rad21l1 plays a role in oogenesis yet is dispensable for spermatogenesis. Moreover, deletion of *rad21l1* activates a Tp53-mediated response in females that does not require the formation of Spo11-dependent double strand breaks. We propose that Rad21l1 functions at a critical step of oogenesis that may provide insight into errors that lead to increased birth defects and miscarriage.

## Results

Mouse *Rad21L* and zebrafish *rad21l1* were identified in silico as a kleisin subunit of cohesin by sequence homology to mouse and human *Rad21* and *Rec8* paralogs [31,45]. To determine if zebrafish Rad21l1 protein is a component of axial elements (AE), lateral elements (LE), and/or the transverse filament (TF) of the synaptonemal complex in zebrafish, we created an antibody to the C-terminus region of Rad21l1 (amino acids 329-516) (S Fig 1). Using this antibody, we stained nuclear surface spread spermatocytes and oocytes using immunofluorescence (IF) detection by 3D-structured illumination microscopy (Fig 1). Previously, we and others showed that the AE protein Sycp3 loads at leptotene near telomeres clustered in the bouquet to form short lines. Shortly thereafter, the axes elongate toward the middle of the chromosome during zygotene until they reach full length at early pachytene [38–41]. Synapsis, as detected by IF staining of the transverse filament protein Sycp1, initiates near the chromosome ends and extends inward, slightly trailing the elongation of the AE until pachytene [39,40]. We found that Rad21l1 foci are dispersed throughout the spread region in leptotene and zygotene yet are concentrated along the unpaired nascent Sycp3 axial elements (Fig 1A [a-h]). At early zygotene, Rad21l1 continues to load on unpaired AE as they elongate, yet as chromosome regions synapse, Rad21l1 is also found between axes and colocalizes with Sycp1 (Fig 1A [m-p]). By late zygotene and pachytene, the dispersed foci largely disappear and nearly all Rad21l1 protein is found along and between synapsed axes (Fig 1A [q-t]). Similar staining pattern is seen in females (Fig 1B). This localization is similar to that described in a recent study using a different antibody to zebrafish Rad21l1 [41]. Localization to axes and the transverse filament of the SC is also similar to the localization of RAD21L protein in mouse as viewed by super-resolution microscopy [46]. These results show that zebrafish Rad21l1 is an abundant protein associated with meiotic chromosome architecture starting at leptotene

**Fig 1.**
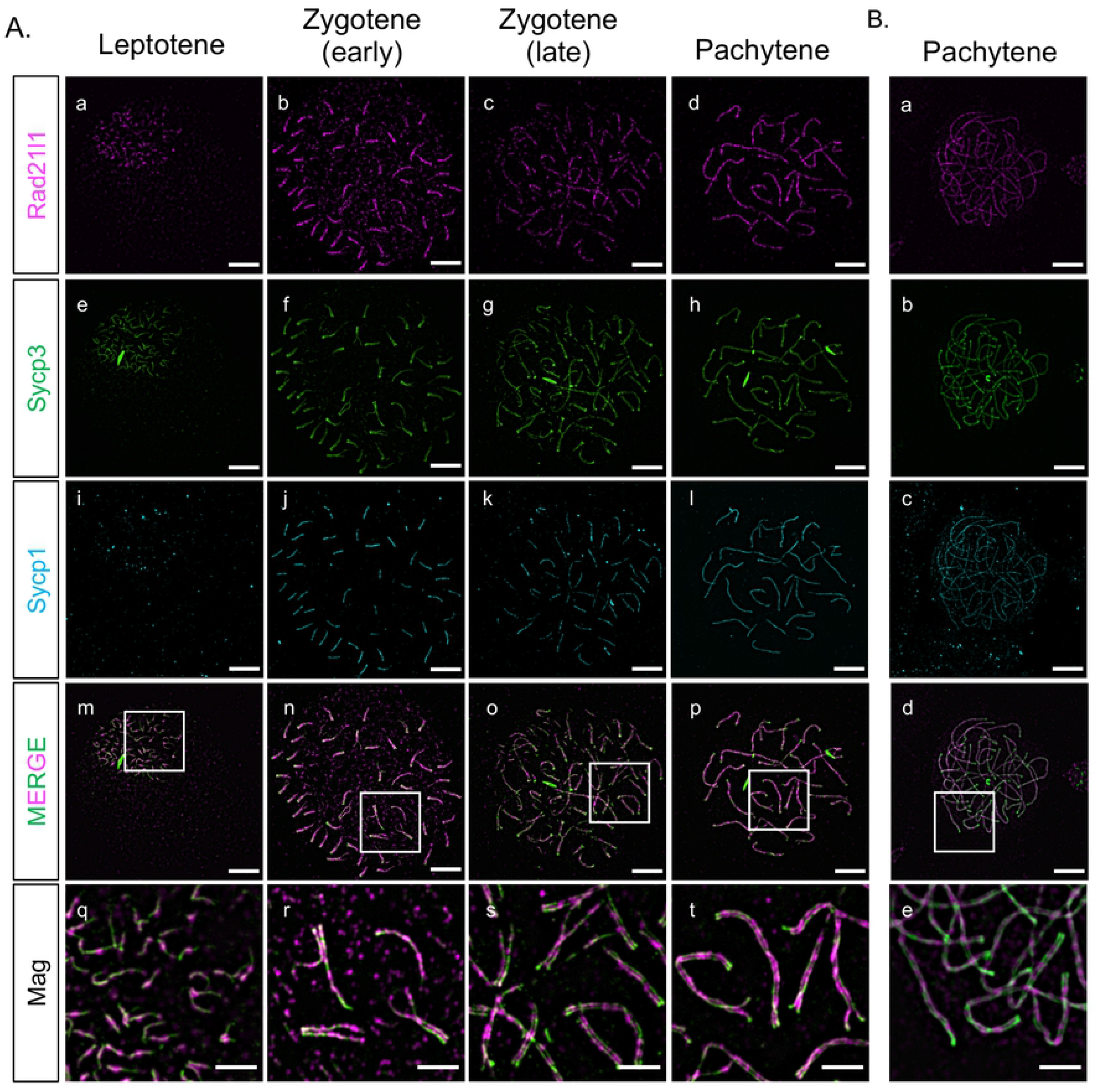
Rad21l1 expression and loading. (A) Rad21l1 loading during prophase I of meiosis in spermatocyte nuclear surface spreads. Rad21l1 (magenta) loads onto chromosome axes simultaneously with Sycp3 (green) and is also dispersed as foci throughout the spread in leptotene. In early zygotene, Sycp1 (cyan) lines start near the telomeres and synapsis extends inward through late zygotene. The merged images are Rad21l1 and Sycp3 channels only. Mag images are magnifications from the Merge panels; the regions magnified are indicated by white boxes. Panel series a-p scale bar = 5 um. Mag panel series q-t scale bar = 2 um. (B) Rad21l1 loading during prophase I of meiosis in oocyte nuclear surface spreads. Panels a-e are arranged similarly to the corresponding panels of part (A).

### Creating the *rad21l1* mutant

We created a *rad21l1* mutant to assess the meiotic function of this cohesin subunit in zebrafish. The *rad21l1* gene in zebrafish consists of 14 exons encoding a 546-amino acid (aa) protein product (NCBI Reference Sequence: NP_001073519.1). We used TALENs targeted to the second exon to introduce an indel mutation by error prone repair, which we designated as the mutant allele *rad21l1^uc89^.* Sequencing of genomic DNA isolated from offspring of founder lines identified a 17 base pair deletion that resulted in a frameshift mutation in the coding region that predicts a truncated protein of 27aa (Fig 2A). To confirm disruption of Rad21l1 expression, we probed spermatocyte nuclear surface spreads from *rad21l1*^*−/−*^ mutants with the anti-Rad21l1 antibody and found that Rad21l1 was absent (Fig 2B). From this we conclude that *rad21l1*^*uc89*^is a null allele (hereafter referred to as *rad21l1^−/−^*).

**Fig 2.**
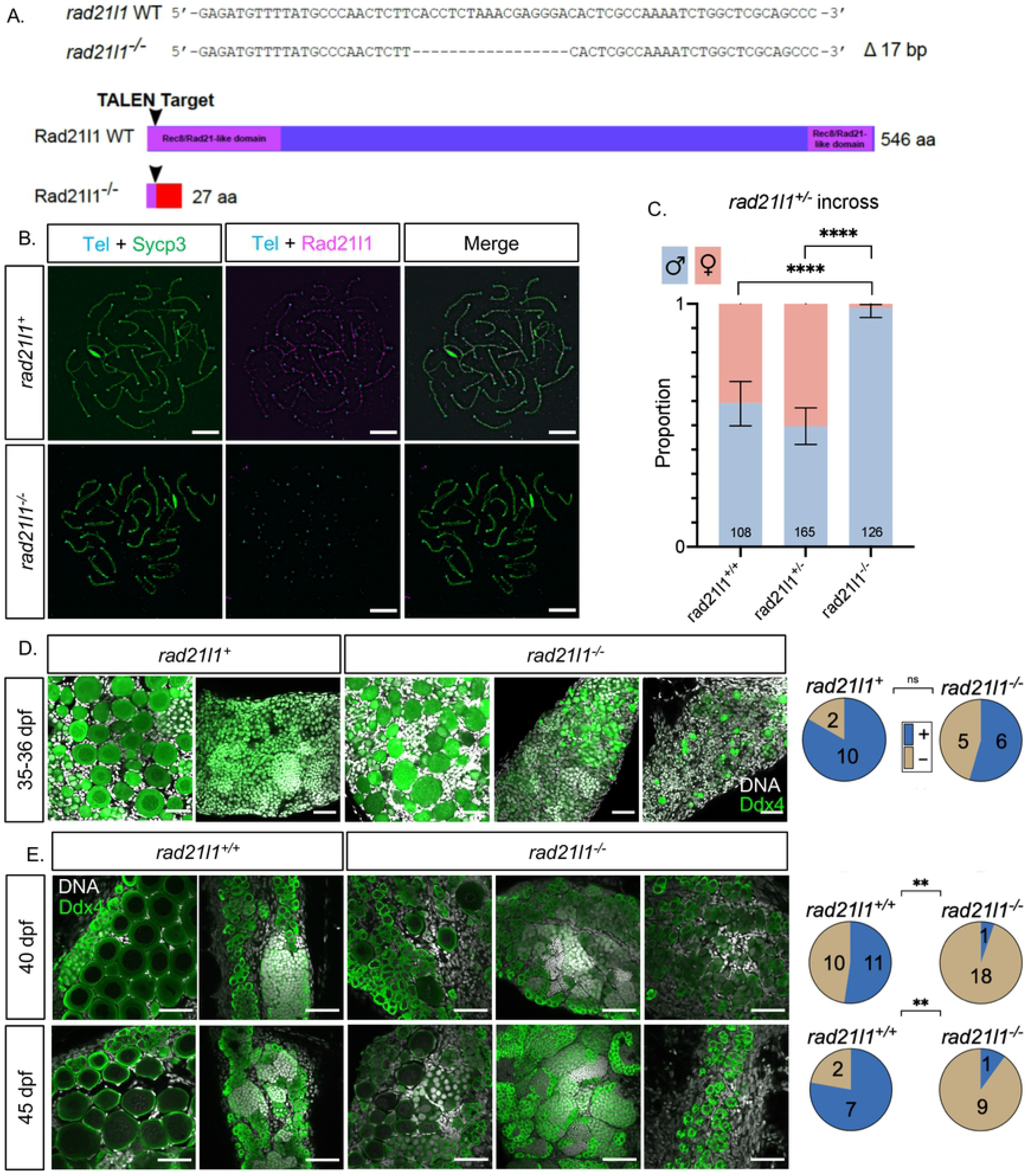
*rad21l1*^*−/−*^ mutants are predominantly male due to late sex reversion. (A) TALEN generated 17-bp deletion leads to a frameshift mutation resulting in a truncated 27 amino acid (aa) Rad21l1 protein with the conserved Rec8/Rad21-like family domains (1-100 aa and 495-543 aa) disrupted or deleted. Rec8/Rad21-like domains (purple boxes); altered amino acid sequence (red box). The ATG translational start site is located at the 4th-6th nt from the end. (B) Spermatocyte nuclear spreads stained for telomeres (cyan), Sycp3 (green), and Rad21l1 (magenta). Rad21l1 forms lines of foci along the Sycp3 axis in *rad21l1*^*+/−*^ spermatocytes. In the *rad21l1* mutant, no lines of Rad21l1 foci are seen. The *rad21l1* mutant spermatocytes can form axes and pair homologs. Scale bar = 5 um. (C) Sexed offspring of a *rad21l1*^*+/−*^ incross show a depletion of females in *rad21l1*^*−/−*^ fish. Data pooled from multiple crosses. (D) Sections of gonads prepared from 35-36 dpf r*ad21l1*^*+/+*^ and *rad21l1*^*+/−*^(labelled *rad21l1^+^)* and *rad21l1*^*−/−*^ fish and stained for DNA (gray) and Ddx4 (also known as Vasa; green). At 35-36 dpf, oocytes are present in 10/12 *rad21l1*^*+/−*^ and 6/11 *rad21l1*^*−/−*^samples. Scale bar = 30 um. (E) Whole mounts of gonads from 40 and 45 dpf are stained for DNA (gray) and Ddx4 (green). At 40 dpf, oocytes are present in 11/21 *rad21l1*^*+/+*^and 1/19 *rad21l1*^*−/−*^ samples. At 45 dpf, oocytes are present in 7/9 *rad21l1*^*+/+*^and 1/10 *rad21l1*^*−/−*^ samples. Scale bar = 30 um. Fisher’s exact test used for all statistical analysis. (+; oocytes present), (−; oocytes absent), ns = p>0.05, ** = p<0.01, **** = p<0.0001.

### *rad21l1* mutants are predominantly male due to late female to male sex reversion

All zebrafish start out with a bipotential gonad that differentiates into an ovary or testis based on a combination of genetic and environmental factors [47–51]. The sex of the gonad is determined by the quantity of oocytes produced during the bipotential phase; females will develop when there are sufficient numbers of oocytes to support ovarian development. In wild-type strains, oocytes in the gonads of presumptive males will begin to apoptose around 20 days post fertilization (dpf) to prepare for testis development, whereas oocytes in the gonads of presumptive females will continue to mature [52,53]. Continued oogenesis is required for zebrafish to maintain the female state; mutants that affect the production of oocytes results in female to male sex reversion [54–56]. We assessed the sex ratio of adult *rad21l1*^*−/−*^ homozygous mutants (n=126), based on protruding belly and morphology of the genital papilla, and found only two females (1.6%), while wild type (n=108) and heterozygous (n=165) fish showed sex ratios within the normal range (40.7% and 50.3% females, respectively; Fig 2C). Interestingly, the two *rad21l1*^*−/−*^ females were able to reproduce and generate healthy offspring, indicating that the sex-reversal phenotype associated with loss of *rad21l1* displays incomplete penetrance. Since sex is determined by both genetic and environmental conditions, it is not known if this failure to sex revert is due to one or the other, or both.

The overabundance of males in the adult mutant population suggests that the *rad21l1*^*−/−*^ mutation may be affecting oogenesis. A precedence for this phenotype is seen in *fancl* and *brca2* mutants where the gonads never form mature oocytes and most animals develop as males [54,55]. To test if *rad21l1*^*−/−*^ mutants can form oocytes, we analyzed gonad sections of 12 animals that were wild-type or heterozygous, and 11 knockout animals at 35-36 dpf stained with DAPI and antibodies to Ddx4, a germ-cell marker also known as Vasa. Here we saw no significant difference in the number of *rad21l1* positive (*rad21l1^*+/+*^or rad21l1^+/−^)* compared to knockout animals with oocytes, suggesting that oogenesis in the *rad21l1*^*−/−*^ mutant can progress through early stages of meiotic prophase (Fig 2D).

To determine the time window at which *rad21l1*^*−/−*^ mutants differentiate as males, we examined gonads of wild-type and *rad21l1*^*−/−*^ animals at 40 and 45 days stained with DAPI and Ddx4. At both time points, wild-type samples could be easily identified as either female or male. That is, 11/21 gonads had oocytes at 40 dpf, and 7/9 gonads had oocytes at 45 dpf (Fig 3E). By contrast, at both 40 and 45 days, the majority of the mutant gonads contained no oocytes (18/19 and 9/10, respectively), indicating a significant decline in oocyte progression in the mutants compared to wild type (p<0.01, Fisher's exact test). In contrast to their wild type tank-mates at 40 and 45 dpf where the gonads already committed to a male or female fate, the mutants exhibited a broad distribution of gonad morphologies, ranging from i) having early stage oocytes, ii) resembling wild-type males, and iii) having primarily premeiotic germ cells (Fig 3E). Since the latter class was not seen among the 35-36 dpf mutants, our interpretation is that this class of gonads represent a delay in testis development following sex reversion (see below). Together, these data suggest that a portion of the *rad21l1*^*−/−*^ males are the product of female to male sex reversion. Notably, in the two cases where oocytes were seen in a mutant gonad, the DAPI signal shows oocytes of different stages of differentiation past pachytene and even up to lampbrush stage where cells have entered diplotene stage (Fig 3E) [57].

**Fig 3.**
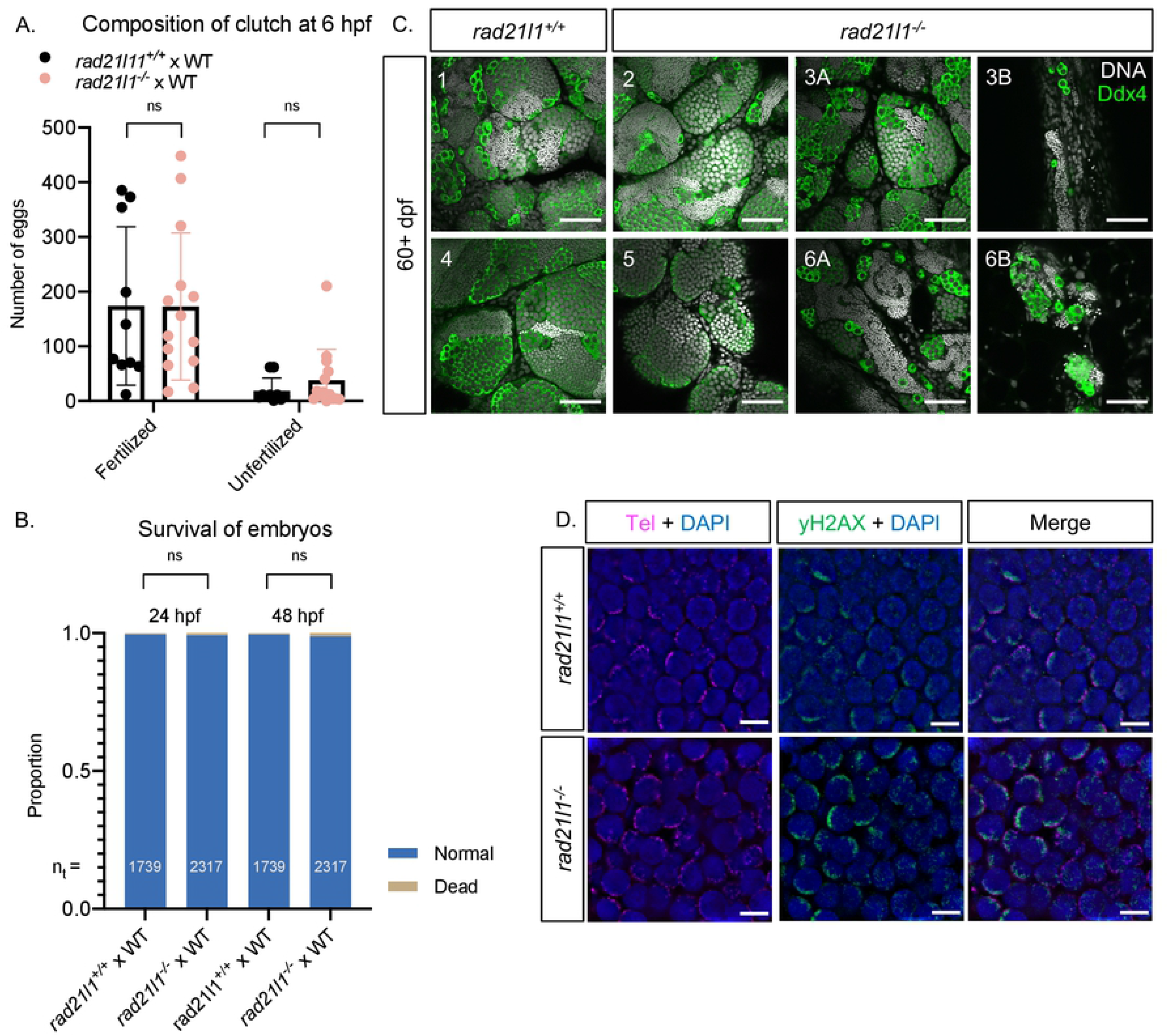
Rad21l1 is dispensable for male fertility. (A/B) Data resulting from test crosses between *rad21l1*^*−/−*^ males and wild-type females to assess fertility and reproductive phenotype. *rad21l1*^*+/+*^male tank mates were used as controls. No significant difference in the number of eggs the males caused the females to release, the composition of the resulting clutch at 6 hpf, or the survival of the embryos through 48 hpf. Data pooled from 14 crosses over 5 weeks using the same pool of 14 *rad21l1*^*−/−*^ males, 12/14 of which crossed successfully at least once. Unpaired, two-tailed student t-test used for statistical analysis, ns = p>0.05. (C) Whole mount adult testes stained for DNA (gray) and Ddx4 (green) showing a phenotypic range of gonad morphology in *rad21l1*^*−/−*^ males. All samples except #5 displayed large clusters of mature sperm. Images marked as A and B were taken from the same sample to show variation within a single gonad. Wild-type tank mates used as controls. Scale bar = 30 um. (D) Testes sections stained with a PNA telomere probe (Tel; magenta), an antibody to _γ_H2AX (green), and DAPI (blue), showing that telomere clustering and DSB localization (_γ_H2AX) are normal in the *rad21l1* mutant. Scale bar = 5 um.

### Rad21l1 is dispensable for male fertility

#### *rad21l1* mutant males produce healthy offspring

To test if the mutant males are fertile, we set up individual crosses using one mutant male and one wild-type female per tank over the course of several weeks (5 crosses each of wild type and mutant males per attempt). While 12 of 14 individual mutant males crossed successfully at least one time, 2 males did not cross, even after three attempts. We next used the pool of 12 fertile mutant males to assess if their progeny exhibit developmental defects. The eggs produced by these crosses were collected and categorized at 6 hours post fertilization (hpf) as either fertilized or unfertilized. There was no significant difference in the fertility of the mutant males compared to their wild-type tank mates (Fig 3A). All normal, fertilized embryos were further incubated at 30°C and observed at 24 and 48 hpf. No significant difference in the frequency of survival or the development of progeny of mutant and wild-type tank mates was observed (Fig 3B).

#### Some *rad21l1−/−* mutant males display unusual gonad morphology

Adult *rad21l1*^*−/−*^ mutant males at 60+ dpf had largely normal-appearing gonads as seen by anti-Ddx4 and DAPI staining of whole mounts, however, the exceptions suggest a depletion of early germ cells during development or delayed development of sex reverted males. Of 11 mutant samples that were stained and imaged, 8 resembled wild-type, 2 had areas of sparse germ cells, and 1 contained no germ cells. One possibility is that mutants that reverted to male especially late were unable to recover an appropriate number of spermatogonia during the late development of testes. Notably, even regions that are sparsely populated by germ cells in the mutant adult gonad have sperm, which supports our finding that the majority of mutant males are fertile (Fig 3C).

#### *rad21l1*^*−/−*^ mutant males are proficient for forming the bouquet as well as pairing and synapsis of homologous chromosomes

In mice, *Rad21l*^*−/−*^ mutants show defects in telomere attachment to the nuclear envelope (Biswas, 2016). We tested if this was the case in zebrafish by staining mounted gonad sections from wild-type and *rad21l1*^*−/−*^ animals with DAPI to detect DNA, a PNA probe to detect telomeres, and an antibody to _γ_H2AX to detect DSBs. In contrast to what was observed in mice, we found that telomere clustering and DSB localization in mutant sections were indistinguishable from wild type (Fig 3C). Additionally, by probing Sycp3 localization on nuclear surface spreads, we found that *rad21l1*^*−/−*^ mutant males form 25 paired bivalents, indicating that Rad21l1 is not essential for these processes in the male (Fig 2B). Together these data suggest that Rad21l1 is dispensable for meiotic progression and fertility in zebrafish males. It is noteworthy that chromosomes in mutant male spreads seemed occasionally to be more fragmented or fragile to the physical forces of the spreading procedure and contained some internally asynapsed regions.

### The *rad21l1*^*−/−*^*;tp53*^*−/−*^ double mutant partially rescues the sex ratio

Previous studies analyzing *brca2* and *fancl* mutants demonstrated that the *tp53* mutation rescues the female to male sex reversal phenotype seen in these mutants, possibly by inactivating a DNA damage checkpoint pathway [54,55]. In mouse spermatocytes, P53 participates in recombination dependent pachytene arrest [58]. To test if the loss of *tp53* could rescue the *rad21l1*^*−/−*^ sex reversal phenotype, we created a *rad21l1*^*−/−*^*;tp53*^*−/−*^ double mutant carrying a loss-of-function *tp53* missense mutation [59]. To isolate this genotype, we incrossed double heterozygous *rad21l1*^*+/−*^*;tp53*^*+/−*^ mutants and sexed the resulting offspring. We found that while *rad21l1*^*−/−*^*;tp53* ^*+/+*^and *rad21l1*^*−/−*^*;tp53*^*+/−*^ mutants produced only the rare female (1/27 and 0/63, respectively), 29% (8/28) of the *rad21l1*^*−/−*^*;tp53*^*−/−*^ double mutants developed as females (Fig 4A). These results suggest that deleting *rad21l1* disrupts oogenesis by activating a Tp53 dependent checkpoint.

**Fig 4.**
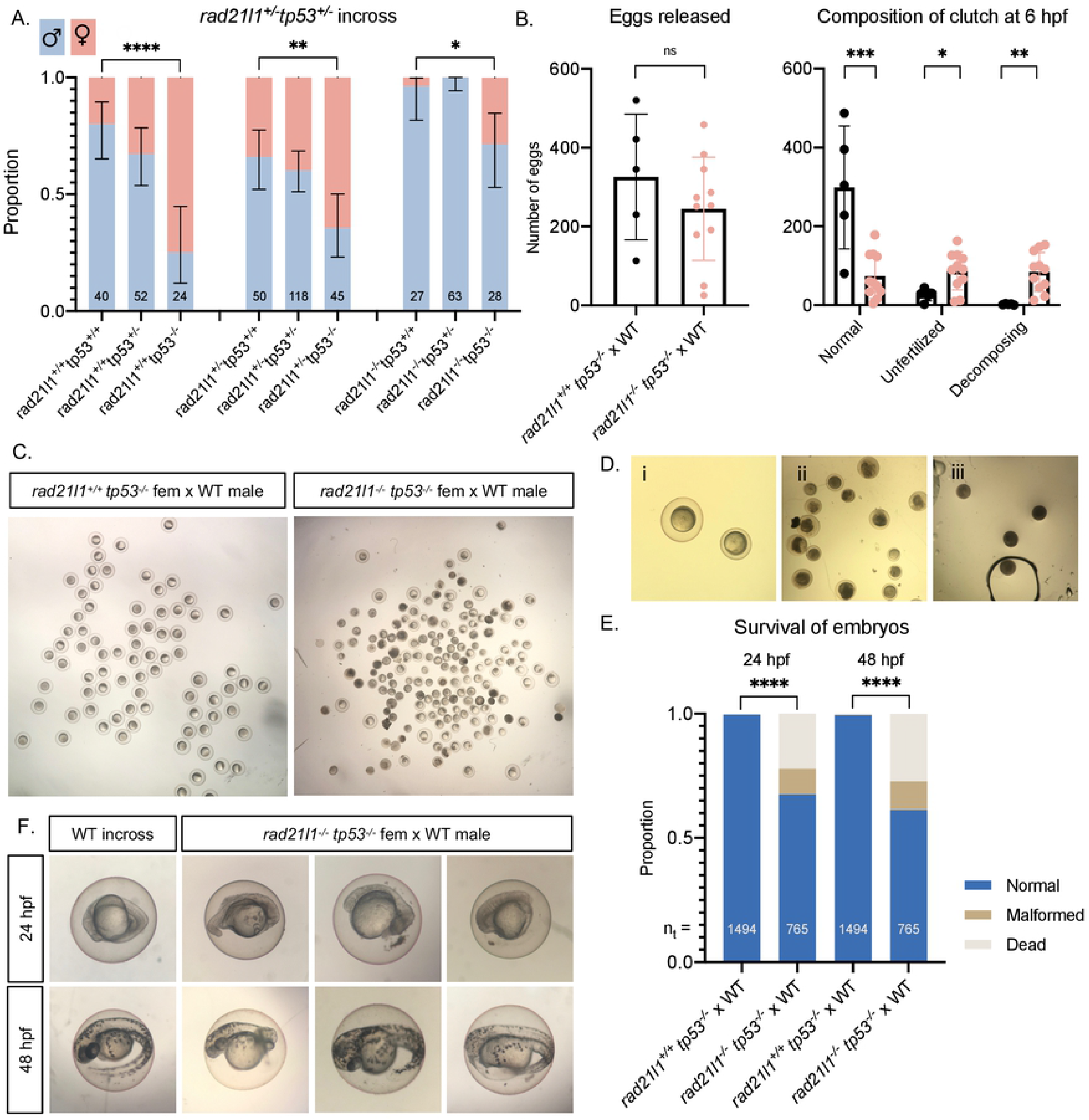
*tp53* knockout restores females to *rad21l1* mutant population, but *rad21l1 tp53* double mutant females produce poor quality eggs and malformed embryos. (A) Sex ratios of all genotypes resulting from a *rad21l1*^*+/−*^ *tp53*^*+/−*^ incross. Data pooled from 3 crosses. Errors bars are 95% confidence intervals. Fisher’s exact test used for statistical analysis. (B) Data resulting from test crosses between *rad21l1*^*−/−*^ *tp53*^*−/−*^ females and wildtype males to assess fertility and reproductive phenotype. *rad21l1*^*+/+*^ *tp53*^*−/−*^ female tank mates used as controls. No significant difference in the number of eggs the females released. *rad21l1*^*−/−*^ *tp53*^*−/−*^ double mutant females release a significantly greater percentage of eggs that fail to be fertilized or display premature decomposition. (C) Representative images of clutches from double mutant and control females at 6 hpf showing lower overall quality of eggs released from double mutant females. (D) Images i-iii show examples of eggs described in the text at 6 hpf. Panel i shows a normal egg (left) and a tiny egg (right). Panel ii shows prematurely decomposing eggs and panel iii shows opaque eggs. All images are the same magnification. (E) Of normal embryos at 6 hpf, 32.4% are dead or malformed at 24 hpf and 38.7% by 48 hpf. Unpaired, two-tailed student t-test used for statistical analysis. (F) Representative images showing the range of malformations seen in developing embryos from *rad21l1*^*−/−*^ *tp53*^*−/−*^ females at 24 and 48 hpf. ns = p>0.05, * = p<0.05, ** = p<0.01, *** = p<0.001.

### *rad21l1*^*−/−*^*;tp53*^*−/−*^ double mutant females produce large numbers of decomposing eggs and deformed offspring

With the *rad21l1*^*−/−*^*;tp53*^*−/−*^ female progeny generated from the double-heterozygous incross, we were able to evaluate the effect deleting *rad21l1* has on oogenesis. If *rad21l1* is dispensable in females, as it is in males, we expected that the double mutant mothers would be fertile and give rise to normal offspring. Alternatively, if the *rad21l1*^*−/−*^ mutation activates a tp53-dependent checkpoint due to errors at some step specific to oogenesis, we expected that the double mutant mothers would produce oocytes, albeit with decreased fertility. To distinguish between these two possibilities, we crossed *rad21l1*^*−/−*^*;tp53*^*−/−*^ females to wild-type males and assessed egg and embryo morphology compared to a *tp53*^*−/−*^ single mutant control. The eggs produced by these crosses were collected and categorized at 6 hours post fertilization (hpf) as normal (fertilized), unfertilized, or decomposing. Fertilized eggs are characterized as being nearly transparent, with a risen chorion and dividing cells. Unfertilized eggs are typically transparent without any signs of decomposition at 6 hpf. Any remaining eggs that had already begun to decompose or did not appear to be correctly formed with a risen chorion were categorized as decomposing.

First, we found that the double mutant females produced high numbers of decomposing eggs and had fewer viable embryos at 6 hpf compared to the *tp53*^*−/−*^ control (Fig 4B-C). Many of the decomposing eggs from the double mutants had a more “opaque” appearance than regular decomposing eggs and did not have a lifted chorion (Fig 4D). This is reminiscent of a previously described phenotype where opaque eggs appeared to be oocytes that failed to progress past stage IV of oogenesis [60]. In addition, many of the eggs from double mutant females were smaller in size and had smaller chorions than normal (Fig 4D). Despite this smaller size, these eggs were considered normal if they fit the criteria described above for the *rad21l1*^*−/−*^ male crosses. We tracked the normal embryos to 24 hpf and 48 hpf (Fig 4E). While the majority of embryos developed normally, we found a spectrum of abnormalities, ranging from normal appearance, to almost a complete failure to develop, to severe head or tail truncations (Fig 4F). These findings suggest that the fertility of *rad21l1*^*−/−*^*;tp53*^*−/−*^ females is severely reduced compared to *tp53*^*−/−*^ controls, with much of the defect arising from poor gamete quality (~⅔), and to a lesser extent the formation of dead and malformed embryos.

Interestingly, in an independent set of crosses we recovered one double mutant female out of 7 that exhibited a normal reproductive phenotype. It is possible that this animal would have developed as a female without sex reversion as was seen for the rare single mutant females (Fig 2C).

### Rad21l1 and Spo11 likely function in different pathways to promote oogenesis

Mammalian oocytes respond to defects in processing DSBs by arresting development and undergoing programmed cell death via a P53 dependent pathway [61,62]. P53 arrest-inducing mutations in the meiosis-specific DSB repair genes *Dmc1* and *Msh5* are suppressed by the elimination of DSBs by deleting *Spo11* [62]. In zebrafish, *spo11* mutants have normal sex ratios and females are fertile, yet give rise to malformed embryos [40,41,63]. We reasoned that if the *rad21l1*^*−/−*^ mutation was inducing meiotic arrest by preventing the repair of Spo11-induced DSBs, eliminating DSBs by deleting Spo11 would rescue the *rad21l1*^*−/−*^ sex reversal phenotype. This was not the case since all of the *rad21l1*^*−/−*^*;spo11*^*−/−*^ double mutants were male (n=17, Fig 5A). This outcome indicates that Spo11 and Rad21l1 act in separate pathways to promote oogenesis. This is further supported by the non-epistatic phenotype of the double mutant as seen in Ddx4 stained whole mounts. The double mutants produce only males as seen in the *rad21l1*^*−/−*^ single mutant yet also fail to produce sperm, as seen in the *spo11*^*−/−*^ single mutants (Fig 5B).

**Fig 5.**
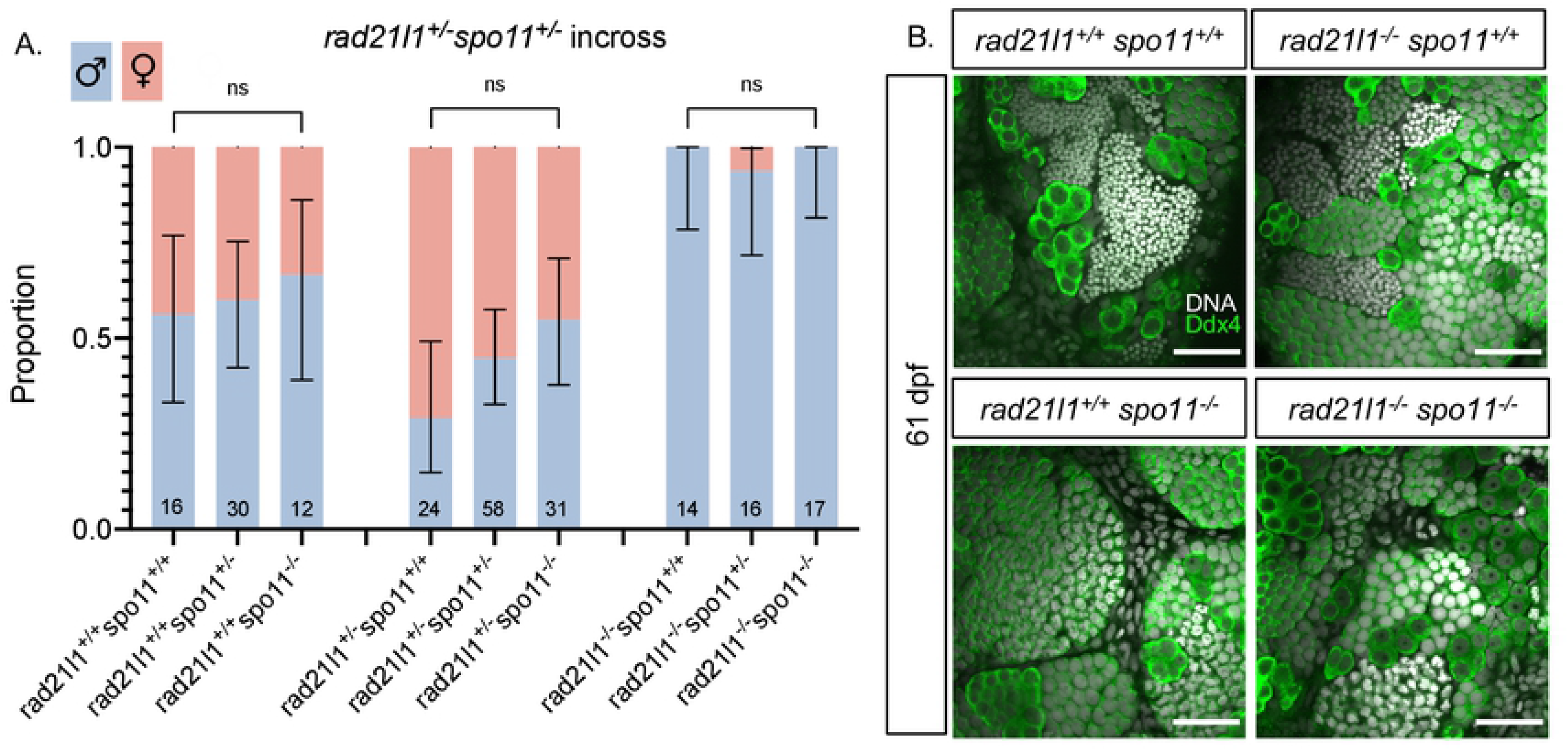
*spo11 rad21l1* double mutants are infertile males. (A) Sex ratios of all genotypes resulting from a *rad21l1*^*+/−*^ *spo11*^*+/−*^ incross. Mostly males seen in all 3 genotypes without Spo11. Data pooled from 6 crosses. Errors bars are 95% confidence intervals. ns = p>0.05. (B) Whole mount testes stained for DNA (gray) and Ddx4 (green). WT and *rad21l1*^*−/−*^*spo11*^*+/+*^samples display clusters of mature sperm, while *rad21l1*^*+/+*^*spo11*^*−/−*^ and *rad21l1*^*−/−*^*spo11*^*−/−*^ samples do not. Scale bar = 30 um.

## Discussion

The data shown here support three major conclusions. First, Rad21l1 is an abundant protein associated with meiotic chromosomes, colocalizing with both the axis protein Sycp3 and the transverse element of the synaptonemal complex Sycp1. Second, spermatogenesis proceeds normally in the absence of Rad21l1 but oogenesis is dramatically affected, as evidenced by female to male sex reversion, poor gamete quality, and an increase in the number of malformed and dead embryos from *rad21l1*^*−/−*^*;tp53*^*−/−*^ double mutant females. Third, the defect imposed by the *rad21l1* mutation results in a Tp53 mediated response that is not seen in males. This response is not due to a failure to form or repair Spo11-induced DSBs. Interestingly, the *rad21l1*^*−/−*^ single mutant females and a *rad21l1*^*−/−*^*;tp53*^*−/−*^ double mutant female displayed normal reproduction, showing that the sex reversal and poor gamete quality phenotypes have reduced penetrance and variable expressivity. We propose that Rad21l1 plays a role in establishing and/or maintaining cohesin integrity that is important for oogenesis but is dispensable in males, and that the loss of Rad21l1 in zebrafish reveals an increased resiliency of spermatogenesis over oogenesis.

While Rad21l1 protein is present on meiotic chromosomes in both males and females, the majority of mutant males were fertile and produced normal progeny. This is in stark contrast to the phenotype of *Rad21l* mutant mice, where defective synapsis leads to arrest at mid-prophase, resulting in azoospermia [31–35,37,45]. This difference could be due, in part, to partial functional redundancy of the meiotic kleisin subunits. While mice have one copy each of *Rec8* and *Rad21l*, zebrafish has two *rec8* paralogs, *rec8a* and *rec8b*, in addition to *rad21l1*. It is possible that one or both of the Rec8 paralogs, or even the mitotic cohesin Rad21, is sufficient to cover for the loss of Rad21l1 in early meiosis in both male and female zebrafish. Some functional redundancy among meiosis-specific cohesins has also been seen in mice [37].

The sexually dimorphic phenotype of the *rad21l1* mutant reveals that oogenesis is more affected by the absence of Rad21l1 than spermatogenesis. The question then arises, what stage of oogenesis is sensitive to the absence of Rad21l1? As in mammals, oogenesis in zebrafish arrests prior to the meiosis I division in the dictyate stage. During arrest, the size of individual oocytes undergoes significant expansion before they are released from the ovary [57]. The expansion includes decompaction of DNA to form lampbrush chromosomes [64]. Thus a time window exists during oogenesis, but not spermatogenesis, where the DNA and/or chromatin architecture may be more vulnerable to damage. In humans, errors in chromosome segregation are a major source of aneuploidy, and the prolonged dictyate arrest is associated with these errors [3,4].

The cause of sex reversion in the *rad21l1* mutant is not entirely clear. Mutations that affect the repair of Spo11-induced DSBs in many organisms lead to check-point mediated arrest of prophase progression [65]. In mouse, the absence of P53 bypasses the pachytene arrest caused by unrepaired Spo11-induced DSBs [58]. If the *rad21l1* mutation prevented the repair of Spo11-induced DSBs, we expected that deletion of *spo11* would also suppress sex reversion; however, this was not observed. Instead, the Tp53-mediated arrest phenotype exhibited by the *rad21l1* mutant may arise from another trigger of the meiotic checkpoint network. One possible explanation is that the mutants experience unrepaired Spo11-independent DSBs. Such breaks could potentially arise from different sources: 1) Oocytes arrested at diplotene may be subject to late DSBs which cannot be efficiently repaired in the absence of Rad21l1, or 2) The absence of Rad21l1 may alter meiotic chromosome architecture that is specific to female meiotic chromosomes in a way that makes them more susceptible to Spo11-independent DSBs.

As oocytes mature, chromosomes undergo significant decompaction, which could make the DNA more vulnerable to breakage. Rad21l1 could be important for establishing the structural and/or spatial context in which spontaneous breaks are repaired during dictyate arrest. In mice, mutation or decreased expression of cohesin subunits confer increased sensitivity to DNA damage [12,66]. In *Drosophila* and yeast, cohesins have been shown to bind to sites of induced DSBs [67,68]. Alternatively, the absence of Rad21l1 itself may increase the sensitivity of chromatin to breakage. For example, tethering DNA loops, involving either sister chromatids or homologous chromosomes could protect DNA from DSBs. In either case, Rad21l1 may be dispensable for spermatogenesis where this sensitive stage of dictyate arrest does not occur.

*RAD21L* variants in humans have been linked to increased maternal nondisjunction of chromosome 21 through GWAS analysis [69], and in mouse a null mutation in *Rad21l* is linked to age-dependent oocyte depletion [32]. While the effect of mutating *Rad21l* in mouse is more severe in males, the nature of this arrest is multifaceted. That is, *Rad21l* mutants arrest at pachytene, likely due, in part, to disruption of the process of meiotic sex chromosome inactivation (MSCI) [32,42]. This is consistent with a single-nucleotide polymorphism in human *RAD21L* linked to azoospermia in Sertoli cell-only syndrome in males [70]. Arrest due to a defect in MSCI would be epistatic to a possible downstream phenotype associated with a *Rad21l* mutation in mouse (i.e. arising in diplotene cells), so determining a later role for *Rad21l* during spermatogenesis remains elusive. Zebrafish, which lack heterogametic sex chromosomes [71], is an excellent model to directly compare sexually dimorphic phenotypes associated with mutations in meiotic genes in males and females since mutant phenotypes can be uncoupled from defects arising from MSCI. Here we show that a *rad21l1* knockout mutation in zebrafish has a much more severe defect in females compared to males, providing additional insight into the molecular basis of the maternal age effect.

## Materials and Methods

### Ethics statement

The UC Davis Institutional Animal Care and Use Committee (IACUC) has approved of this work under the protocol #20199; For noninvasive procedures (e.g. fin clips for genotyping), zebrafish were anesthetized using tricaine. Invasive surgical methods were performed on fish euthanized by submerging fish in ice water.

### Zebrafish strains

Zebrafish husbandry was performed as previously described [72]. The wild type NHGRI strain was used in the production of the *rad21l1*^*uc89*^mutants. Fish used in experiments were outcrossed to the AB strain background 3-4 times. The *spo11*^*−/−*^ strain is in the AB background and described in Blokhina 2019. The *tp53*^*−/−*^ mutant is described in [59]. All test crosses were performed with wild type AB strain fish.

### *rad21l1*^*−/−*^ mutant generation

The *rad21l1*^*uc89*^ mutants were generated using transcription activator-like effector nucleases (TALENs) to target exon 2 and genotyped using high resolution melt analysis (HRMA). TALEN target sequences: NG-NI-NG-NH-HD-HD-HD-NI-NI-HD-NG-HD-NG-NG-HD-NI-HD-HD-half repeat NG and NH-HD-NH-NI-NH-HD-HD-NI-NH-NI-NG-NG-NG-NG-NH-NH-HD-NH-half repeat NI. Injected founder fish were raised to adulthood and outcrossed to wild type fish. The resulting offspring were screened for mutations in *rad21l1* via HRMA and subsequent sequencing. HRMA primer sequences are: Fwd 5’-CGCCGAGACATGTTTTATGCCC-3’, Rev 5’-TCAAACACGTGGGCTTTGGT-3’. The HRMA was performed with 20X Eva Green dye (VWR, Radnor, PA, Catalog #89138-982) using a CFX-96 real time PCR machine and Precision Melt Analysis software (BioRad, Hercules, CA). Mutants were backcrossed to either AB or NHGRI strain. The sex reversal phenotype was specific to populations genotyped as *rad21l1*^*−/−*^ indicating that it is unlikely due to off-target effects. The phenotype correlation remained consistent through 5-6 crosses.

### Genotyping

#### Mutant identification

Genomic DNA was extracted and samples were analyzed with HRMA [40]. Primers for *Rad21l1* genotyping were the same as described in the *rad21l1* mutant generation. Primers for *Spo11* were Fwd 5’-TCACAGCCAGGATGTTTTGA -3’ and Rev 5’-CACCTGACATTGCAGCA-3’ with an annealing temperature of 61° C. Primers for Tp53 were Fwd 5’-CTCCTGAGTCTCCAGAGTGATGA-3’ and Rev 5’-ACTACATGTGCAATAGCAGCTGC-3’. Genomic DNA was extracted and samples were analyzed as described for *rad21l1* mutants except that the reaction was done in 2 mM MgCl_2_ with an annealing temperature of 65° C. Two HRMA runs were required to confirm the three genotypes resulting from a *tp53*^*+/−*^ incross; the first run distinguished heterozygous from homozygous samples. Homozygous samples were run again under the same conditions but spiked with wild-type DNA in order to differentiate wild-type and mutant samples.

### Antibody generation

Guinea pig anti-zebrafish Rad21l1 polyclonal antibody production: An N-terminal fragment of Rad21l1 cDNA was amplified with Phusion DNA polymerase (Thermo Fisher Scientific, Catalog #: M0530L) using the following primers: Fwd 5’-aactttaagaaggagatataccatgTCAAGCTTTTGCCTTCCTGT-3’ and Rev 5’-tctcagtggtggtggtggtggtgctcAAGCATGCAGAAAAATAAGGCT-3’. The Rad21l1 PCR product was then cloned into pET28b using NEBuilder HiFi DNA Assembly Master Mix (NEB, Catalog #: E5520S). BL21 (DE3) cells containing pRARE and Rad21l1 overexpression construct were grown in 2.6 L of LB with kanamycin and chloramphenicol until an OD600 = 1 and induced with a final concentration of 1 mM IPTG at room temperature for six hours. The Rad21l1 peptide was purified under denaturing conditions using Novagen NiNTA purification resins (Sigma, Catalog #: 70666) according to the manufacturer’s instructions. The Rad21l1 peptide was concentrated to a final concentration of 1 mg/ml in PBS using a 10 kDa centrifugal filter (Sigma, Catalog # UFC901008). The Rad21l1-derived peptide was injected into three guinea pigs by Pocono Rabbit Farm and Laboratory following the 91-day polyclonal antibody production protocol.

### Chromosome spreads and staining

All chromosome spreads and staining were performed as previously described [40,73]. Antibodies and dilutions described in S1 Table.

### Adult testis section and whole mount preparation and staining

Protocols including “whole mount testes staining “ and “testes section preparation and staining” were performed as previously described [40]. Antibodies and dilutions described in S1 Table.

### Whole mount juvenile gonad staining

Juvenile gonad staining was performed similarly to the adult protocol with some modifications:

#### Dissection and fixation

Euthanized fish were decapitated and cut open along the ventral midline to expose the body cavity. Alternatively, an additional cut was made at the anal fin to expose body cavity if fish was too small to make a ventral cut. The fish were fixed in 4% PFA in PBT at 4° C for 16-18 hours with gentle rocking. The fish were placed into fresh tubes and washed in 0.2% PBT 3 times for a minimum of 5 minutes each. Gonads were dissected out in PBT and placed into a ceramic 12-well plate, 1 gonad per well.

#### Primary antibody staining

Gonads were washed in an antibody block composed of 5% goat serum and 5% BSA in 0.2% PBT for 1 hour minimum on a 2D rocker at room temperature. Primary antibody chicken anti-Ddx4 [40] was added at 1:500 final dilution. Plate was left rocking gently overnight at 4° C.

#### Secondary antibody staining

The gonads were washed 3 times for a minimum of 30 minutes in PBT, then washed in antibody block as described above. Secondary antibody anti-chicken Alexa Fluor 488 was added at 1:300 final dilution. Plate was left rocking gently overnight at 4° C.

#### Glycerol dehydration and mounting

The gonads were washed 2 times for a minimum of 10 minutes each and dehydrated in a series of glycerol (Sigma-Aldrich, Catalog #: G5516-1L) washes for 1 hour minimum each: 30% glycerol with DAPI at 1:5000 dilution in PBT, 50% glycerol with DAPI at 1:5000 dilution in PBT, and 70% glycerol in PBT without DAPI. The gonads were mounted in 70% glycerol without DAPI on slides with vacuum grease applied to the four corners to hold the coverslip in place. Slides were stored at 4° C until imaging.

### Imaging

All images were collected at the Department of Molecular and Cellular Biology Light Microscopy Imaging Facility at UC Davis. Chromosomes spreads were imaged using the Nikon N-SIM Super-Resolution microscope in 3D-SIM imaging mode with APO TIRF 100X oil lens. The images were collected and reconstructed using the NIS-Elements Imaging Software. Sections and fluorescent whole mounts were imaged using the Olympus FV1000 laser scanning confocal microscope. Images were processed using Fiji ImageJ software. Only linear modifications to brightness and contrast of whole images were applied. Images of eggs and embryos were acquired on a dissecting microscope.

### Test crosses

To analyze fertility, individual mutant fish were placed in a divided mating tank overnight with a single AB strain wild type fish of the opposite sex. The divider was removed soon after onset of light, and any eggs produced were collected with a strainer, rinsed thoroughly with system water, and placed in a petri dish at 30° C. At 6 hours post fertilization (hpf), embryos were transferred to embryo medium (1X E3 media has final concentrations of 5 mM NaCl, 0.17 mM KCl, 0.3 mM CaCl_2_ dihydrate, 0.33 mM MgSO_4_ heptahydrate, 6 uM methylene blue) and categorized. Fertilized eggs were kept at 30° C and monitored at 24 and 48 hpf for morbidity and mortality.

## Acknowledgements

We thank James Amatruda for the gift of the antibody to _γ_H2AX and Trent Newman, Kelly Komachi, Ivan Olaya, and Roberto Pezza for discussions.

---

**S1 Fig. Alignment of zebrafish Rad21l1, Rec8a, and Rec8b proteins**. Alignment of zebrafish Rad21l1 (ENSDARP00000074083), Rec8a (ENSDARP00000116796), and Rec8b (ENSDARP00000091417) using the Snapgene (v 5.1.4.1) Clustal Omega tool. Yellow shading indicates amino acids of Rec8a and Rec8b that match the Rad21l1 references sequences. The consensus sequence threshold has set at > 50%. Amino acids 329-516 (highlighted) were expressed to create the Rad21l1 antibody in Guinea pigs.

**S1 Table. Antibodies used in this study**

**S1 File. Master data sheet**

